# Fighting over defence chemicals disrupts mating behaviour

**DOI:** 10.1101/2021.02.12.430958

**Authors:** Sarah Catherine Paul, Caroline Müller

## Abstract

Studies on intraspecific contest behaviour predominantly focus on contests between individuals of the same sex, however contest behaviour is also expected to occur between individuals of the opposite sex including possible mates. Here we investigate potential trade-offs between mating and fighting behaviour in the turnip sawfly (*Athalia rosae*). Adults of this species collect chemical defence compounds (clerodanoids) directly from plants but also indirectly by nibbling on conspecifics that have already obtained clerodanoids themselves, a highly aggressive behavioural interaction. An *A. rosae* individual without clerodanoids may therefore be the potential mate or attacker of an individual of the opposite sex that has gained clerodanoids. We paired males and females with or without clerodanoid access and manipulated body mass differences between the sexes via the early life starvation of females. We show that asymmetrical clerodanoid acquisition between male-female pairs causes an increase in agonistic nibbling behaviour, irrespective of sex. Moreover, fighting over clerodanoids disrupted mating behaviour, and the frequency of aggressive nibbling behaviour in these pairs was determined by the comparative body mass of the attacking individual. Our study highlights the vital importance of investigating agonistic intersex interactions not only over mating but also over resources.

## Introduction

Conflict between individuals of the same species is part of the fabric of animal lives, shaping their life histories and influencing fitness [1]. These agonistic interactions occur over resources that are limited or which vary in quality, such as food [2], shelter [3], oviposition sites [4], and members of the opposite sex (both male [5] and female [6]). Whether a fight occurs depends on the costs (C) of engaging in a fight and the fitness benefits, or value (V), of winning the resource [7,8]. Contest outcome is determined by the interplay between the physical capability of an individual to win a fight, its so called resource holding potential (RHP), and the motivation of an individual to win a fight based on the perceived and actual quality of a resource, known as the resource value (RV) [4,9]. The large majority of research on intraspecific dyadic contests focuses on conflict between individuals of the same sex [10,11], however a raft of intersex interactions also occur in nature [e.g. 12]. Such contests may add an additional dimension to the matrix of factors influencing not only an individual’s decision to fight but fight outcome. For example, an opponent with a resource may at the same time be a potential mate, setting up an interesting trade-off between mating and fighting, but such potential trade-offs remain little explored.

In addition to the usual costs of fighting, including energetic costs [13] and increased predation risk [14], agonistic intersex interactions may result in the loss or reduced likelihood of mating with the opposing individual. Mating is likely to have the most immediate fitness benefit [15] unless the resource which could be gained via intersex fighting increases future reproductive success, for instance by winning a territory [16,17]. In such circumstances the fitness benefits of immediate mating might be outweighed by the benefits of increased future reproductive success [18,19]. Such a scenario is of particular relevance to systems in which mating opportunities are abundant and both sexes mate multiple times, as is common in many insect species [10,20].

Depending on the mode of assessment, factors determining the likelihood of winning a fight (i.e. RV and RHP) could also feedback into whether a contest escalates to involve a contact fight in intersex contests [21]. For example, high quality individuals may not benefit significantly from gaining a resource, lowering their intrinsic RV [22] and shifting the balance of costs and benefits to favour mating over fighting. Furthermore, even if an individual would benefit from obtaining the resource (high RV) it may not necessarily be able to win a contest if it has a lower RHP than its opponent [23, but see also 24]. This is particularly relevant for intersex interactions, as sexual dimorphism often leads to significant differences between the sexes in body size [25] and thus RHP [26]. Moreover, the strong link between RHP and condition [27,28] means that a low RHP in one individual might not only be linked to sexual dimorphism but also to poor mate quality [29] with consequences for mate choice [30,31]. It may therefore be more advantageous to mate with an individual that has a higher RHP than to attempt to engage in contest behaviour for a resource. Thus, a number of different and potentially interacting factors may contribute to the outcome of intersex interactions between potential contest and mating partners.

Here we investigated intersex interactions in the turnip sawfly *Athalia rosae* (Hymenoptera: Tenthredinidae). Adults of *A. rosae* collect clerodane diterpene compounds, from now on called clerodanoids, from non-food plants [32]. Clerodanoids act as a deterrent to several predators of *A. rosae* [33] and as such represent a significant resource. Adults fight each other to gain access to these resources, as they can gather clerodanoids via aggressively nibbling on the exterior of conspecifics [preprint 34]. This behaviour not only carries the risk of lowering the defender’s chemical defence levels, but may increase its vulnerability to predation occurring during the agonistic interaction itself, as agonistic behaviours are known to increase predation risk [35] by reducing vigilance behaviour [14]. The mating success of *A. rosae* females, but not that of males, has also been shown to be increased by the possession of clerodanoids [36], indicating a potential role of clerodanoids as a female sex pheromone. The use of sex pheromones, including those derived from plant secondary compounds being common in insects [37]. Due to the fact that female *A. rosae* are heavier and larger than males [38,39] this difference in mating success may be driven by fighting behaviour between individuals with or without clerodanoid access, where males with clerodanoids (C+) are unable to mate with females without clerodanoids (C-), because the latter are able to aggressively nibble on C+ males due to their higher RHP.

By recording both the mating and fighting behaviour of *A. rosae* male-female pairs with either symmetrical or asymmetrical clerodanoid acquisition (i.e. ♀C+♂C+, ♀C+♂C-, ♀C-♂C+, ♀C-♂C-) we aimed to establish the degree to which these behaviours are influenced by clerodanoids. We predicted that both mating and agonistic behaviour would be most strongly influenced by female clerodanoid status. When females had clerodanoids (C+) mating would occur more quickly and agonistic behaviour (e.g. fighting or aggressive nibbling) would be less frequent than in the control (♀C-♂C-) and that the reverse would occur when females did not have clerodanoids but males did (♀C-♂C+). In addition, we tested the role of body mass in determining the outcome of the asymmetrical interactions, through its effect on RHP, by using females that were starved during larval development [(S)♀C+♂C-], which is known to lower adult mass in *A. rosae* [40]. We predicted that lower female body mass would result in an increase in fighting and a decrease in mating behaviour in comparison to the control.

## Methods

### Experimental rearing of sawflies (F0-F2)

Sixty adults of *A. rosae* (F0) were collected in a meadow in Verl, Germany (51°52′23.0″N 8°33′32.0″E) in July 2018 and split into two breeding groups, A and B, placed each in a mesh cage (60 x 60 x 60 cm). Females in each cage were provided with plants of *Sinapis alba* (Brassicaceae) for egg laying and emerging F1 larvae were provided with plants of *Brassica rapa* var. *pekinensis* (Brassicaceae) as food. Larvae of the last instar (eonypmhs) were placed in individual pots containing ~30 g sterilised soil for pupation and after emergence F1 females were mated with an F1 male from the opposite breeding group (e.g. F1♀A x F1♂B). Post mating each mated F1 female (N=30, produce male and female offspring) and each virgin female (N=30, produce only male offspring [41]) were given an individual S. *alba* plant for oviposition and provided *ad libitum* with a honey–water mixture (1:50). Emerging F2 larvae were collected daily from each plant and the larvae from each female were split evenly between two ventilated containers (25 cm x 15 cm x 10 cm), one for each larval starvation condition [no starvation and starvation (S)]. Larvae were provided with *ad libitum* middle-aged *B. rapa* leaves and moistened tissue paper to prevent desiccation, except larvae of the (S) condition, from which every third day all *B. rapa* leaves were removed for 24 hours. Eonymphs were placed individually in soil pots for pupation and after emergence F2 adults were placed in individual Petri dishes (35 mm) with honey water-infused tissue paper as a food supply. These adults were kept at ~5 °C in a refrigerator until use in behavioural assays, which due to their short life span [38] prolonged the period over which the experimental work could be carried out. All rearing was carried out in a climate chamber (temp: 20 °C:16 °C (16 h:8 h), light:dark (16 h:8 h), 70 % r.h.) and all host plants were grown from seeds in a greenhouse (no climate control, light:dark 16 h:8 h).

### Behavioural assays

Adult *A. rosae* (3-18 days post eclosion) were removed from the refrigerator 48 hours prior to the commencement of the behavioural assays. In order to overcome any difference in the behaviour of the first mating of virgin males [42], males were mated to a non-focal female without access to clerodanoids 48 h prior to their focal mating assay. Then, all adults were weighed to the nearest 0.01 mg (Sartorius AZ64, M-POWER Series Analytical Balance, Germany) and provided with a honey-water mixture. C+ individuals were additionally provided with a small section (1 cm^2^) of a leaf of *Ajuga reptans* (Lamiaceae) for 48 h, giving them the opportunity to incorporate clerodanoids prior to the start of the trial. Plants of *A. reptans* were collected from a population at the edge of a local forest (52°01′58.2″N 8°29′04.5″E) in the summer of 2018. This plant species does not serve as food plant but is used by *A. rosae* to gather clerodanoids [32]. Individuals that either were or were not exposed to *A. reptans* (i.e. C+ or C-) were handled with different forceps and forceps were cleaned with 70 % ethanol in between each use to prevent an inadvertent transfer of chemical compounds between individuals.

Behavioural assays were carried out between male and female *A. rosae* across five different treatment levels (♀C+♂C+, ♀C+♂C-, (S)♀C+♂C-, ♀C-♂C+, ♀C-♂C-; figure 1) set up to investigate the effects of clerodanoid access and female early life starvation on mating and agonistic behaviours (table 1). For each of the five treatment levels, an individual female was first added to a mating arena, consisting of a Petri dish (60 mm × 15 mm), and one male was then added to the opposite side of the arena (N = 11 to 18 replicates per treatment level, figure 1). Interactions were recorded for 25 min or until copulation had finished, using a Sony HDR-CX410VE (SONY EUROPE B.V.) camcorder (AVCHD - 1920 x 1080 - 25 fps). Video data was analysed blind by the same observer using the software BORIS v 7.9.8 [43]. All recorded agonistic contact and mating behaviour was observed at 0.3x the original speed and at 2x display magnification to ensure that each behaviour was categorised and timed correctly.

**Figure 1.**
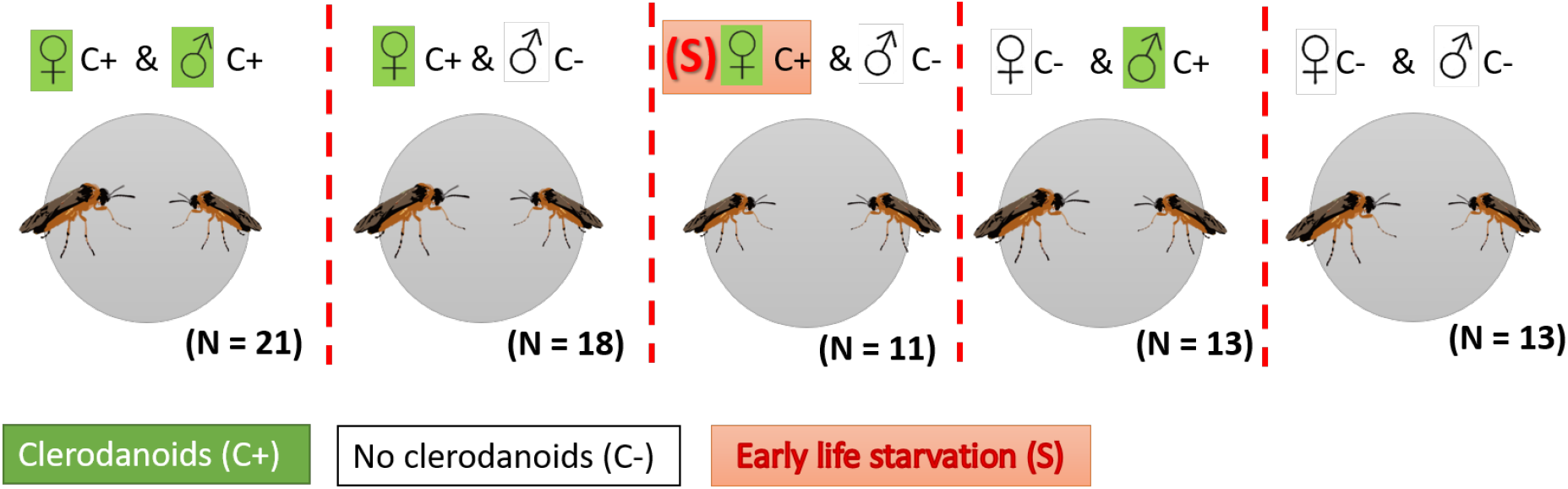
Five different treatment levels and numbers (N) of individual pairs of *Athalia rosae* used in the behavioural assays. Green boxes denote those individuals who had access to clerodanoids via nibbling on *Ajuga reptans* leaves (C+), those without green boxes had no access to *A. reptans* leaves and therefore do not have clerodanoids (C-), and the red box denotes early life starvation (S).

**Table 1.** Behaviours recorded during mating trials of *Athalia rosae* with descriptions of each behaviour. Behaviours are divided into agonistic behaviour and mating behaviour. State behaviours have their occurrence and duration recorded and point behaviours their occurrence but not duration (see S1 for example video).

| Behavioural code | Behavioural type | Description |
| --- | --- | --- |
| **Agonistic contact behaviour** |  |  |
| **Front limb battling** | State | Both individuals face each other and use front legs in 'battling' motion. Front legs make contact and both individuals engage in behaviour. |
| **Fighting / attempted nibbling** | State | Individuals use all legs and move entire body when fighting. Distinguished from nibbling by a lack of contact of mouthparts with the body of the other individual. |
| **Successful nibbling** | State | Mouthparts of attacker make contact with the body of the defender – individual-specific behaviour. Duration of nibbling for each individual based on the criteria of mouthpart contact. |
| **Agonistic behaviour prior to copulation** | Point | 'Successful nibbling' and/or 'Fighting/attempted nibbling' (i.e. an antagonistic interaction) before mating was classified as any of these behaviours that started before mating occurred; it did not have to finish before mating also occurred. |
| **Agonistic behaviour ends copulation** | Point | 'Successful nibbling' and/or 'Fighting/attempted nibbling' (i.e. an antagonistic interaction) starting before the end of copulation and ending after the end of copulation. |
| **Mating behaviour** |  |  |
| **Attempted copulation** | Point | Male moves/flies towards female with curved abdomen, but with no attempted nibbling (movement of head towards thorax/abdomen/wings/head). |
| **Copulation** | State | Copulation initiated. Start of copulation coded as the moment that the genitals become attached (male inserts genitalia into female). When mating occurs individuals often become still, facing in opposite directions with their wings erect, however if they are fighting at the start of copulation this does not always happen immediately, therefore though this posture is indicative of mating, it does not necessarily indicate the correct start of copulation. |
| **Walking during copulation** | State | Female-specific behaviour. Classified as the movement of a female during copulation, for a distance of at least half a body length in a forward direction. |

### Statistical analysis

All data were analysed using R version 4.0.2 (R Core Team, 2020-06-22). Alpha level was set at 0.05 for all tests and model residuals were checked for normality and variance homogeneity. Models of count data were all tested for both zero-inflation [44] and overdispersion, and models were chosen accordingly. Likelihood ratio tests were employed to establish significance of main treatment effect. Posthoc analyses were carried out using ‘emmeans’ v.1.5.0 [45] for beta regression models and ‘multcomp’ v. 1.4-13 [46] for all other models; treatment levels with clerodanoids were compared to C-C-as the control (Tukey HSD), with the exception of female mass and male-female mass difference (female mass – male mass) as the apriori expectations were different. The effect of starvation during larval development on female mass and the consequent mass difference between males and females was assessed using a linear model (LM) (package: ‘MASS’ v 7.3-51.6), in which treatment was the predictor variable and female mass and mass difference were the response variables for each model. Variation in whether copulation occurred was assessed using a binomial model (package: ‘MASS’) with copulation occurrence as the response variable and treatment as the predictor variable. The effect of treatment on the time taken for copulation to commence and copulation duration was assessed using an LM (package: ‘MASS’), in which either log[time to copulation (s)] or copulation duration (s) were the response variables and treatment was the predictor variable in both cases. The influence of treatment on the number of failed mating attempts was tested using two models in order to account for both separation effects (0 and 1 all in one treatment level) and overdispersion. A binomial model, fitted using maximum penalised likelihood to handle separation [package:‘brglm’ v. 0.7.1;, 47], was used to assess whether there was a difference between treatment levels in whether failed/attempted copulations occurred. A negative binomial model (glm.nb, package: ‘MASS’) was used to assess whether the number of failed/attempted copulations per pair (response variable) varied between treatment levels (predictor variable), with log[assay duration (s)] fitted as an offset to account for differing assay durations. The effect of treatment on whether females walked during copulation and on the proportion of time that females spent walking during copulation was modelled using a binomial model, fitted using maximum penalised likelihood (package:’brglm’), and beta regression [package:’betareg’ v.3.1-3;, 48], respectively.

Variation in the occurrence of leg battling, fighting/attempted nibbling, successful nibbling, agonistic interactions pre-copulation or across the end of copulation (i.e. copulation interrupted by agonistic interaction), across treatment levels (predictor) was assessed using individual binomial models, fitted using maximum penalised likelihood to handle separation when it occurred (packages: ‘MASS’ or ‘brglm’). The effect of treatment on the rate (occurrence per second) of leg battling, fighting/attempted nibbling, or successful nibbling was assessed using individual negative binomial glms (glm.nb, package: MASS). Each of these three response variables was modelled separately with treatment as the predictor variable and log[assay duration (s)] as an offset. How the proportion of time during the assay that females spent engaging in fighting/attempted nibbling, and total successful nibbling varied depending on treatment (predictor) was modelled using individual beta regressions (package: ‘betareg’). Because the occurrence of successful nibbling differed between treatment levels, a further analysis was carried out to establish the identity of the successful nibblers within a pair using a binomial model (package: ‘MASS’), where nibbling occurrence was the response variable and clerodanoid exposure, sex, their interaction, and pair ID were the predictors. PairID was set as a fixed opposed to random effect to avoid model overfitting. It is worth noting at this point that unlike contest interactions, in which there is one winner or loser, here the successful nibbling of one individual on another does not preclude reciprocal nibbling from the other individual in a pair. Thus, analyses taking into account both individuals in a pair is valid [49]. The effect of agonistic behaviour on the time until copulation was modelled using an LM with log[time until copulation (s)] as the response variable and the occurrence of precopulatory agonistic behaviour as the predictor. Whether copulation duration (s) was affected by agonistic behaviour was also modelled using an LM with copulation duration (s) as the response variable and occurrence of agonistic behaviour at the end of copulation as the predictor.

## Results

Females that were starved during development had a significantly lower body mass (F_4,62_ = 10.69, p < 0.001, figure 2A) and therefore also lower mass difference to the males they were paired with (F_4,62_ = 6.31, p < 0.001, figure 2B) than females who were not starved during development, across all treatment levels (S2). There was no significant effect of treatment on the number of pairs in which successful copulation occurred (X^2^_4,62_ = 7.89, p = 0.096; % of pairs that copulated: ♀C+♂C+ = 92%; ♀C+♂C- = 72%; (S)♀C+♂C- = 73%; ♀C-♂C+ = 46%; ♀C-♂C-= 54%). For those pairs in which successful copulation occurred there was no significant effect of treatment on copulation duration (F_4,40_ = 1.56, p = 0.205). There was no difference between treatment levels in whether failed copulation attempts occurred (X^2^_4,62_ = 6.77, p = 0.149). However, in those treatment levels where failed copulations did occur the number of failed copulation attempts differed significantly based on treatment (X^2^_4,8_ = 13.28, p = 0.004), a result driven by the higher number of failed mating attempts in the ♀C-♂C+ treatment level compared to the control (♀C-♂C+ vs ♀C-♂C-: estimate (+SE) = 1.61 (+0.67), z = 2.38, p = 0.045). Time until copulation commenced was significantly affected by treatment (F_4,40_ = 2.77, p = 0.040; figure 2C), but although pairs in the ♀C+♂C+ treatment level appeared to have mated more quickly than those in the control, this difference was non-significant (♀C+♂C+ vs ♀C- ♂C-: estimate (+SE) = −1.59 (+0.66), z = −2.40, p = 0.053; figure 2C). There was a significant effect of treatment on whether or not females walked during copulation (X^2^_4,40_ = 9.71, p = 0.046) and on the proportion of time females spent walking during copulation (Pseudo-R^2^ = 0.337; (LRT) X^2^ = 15.15, df = 4, p = 0.004), the latter driven by females in the ♀C-♂C+ treatment level who walked for a longer proportion of the copulation duration than those in the control (♀C-♂C+ vs ♀C-♂C-: estimate (+SE) = 0.20 (+0.05), z = −3.65, p = 0.002; figure 2D).

**Figure 2.**
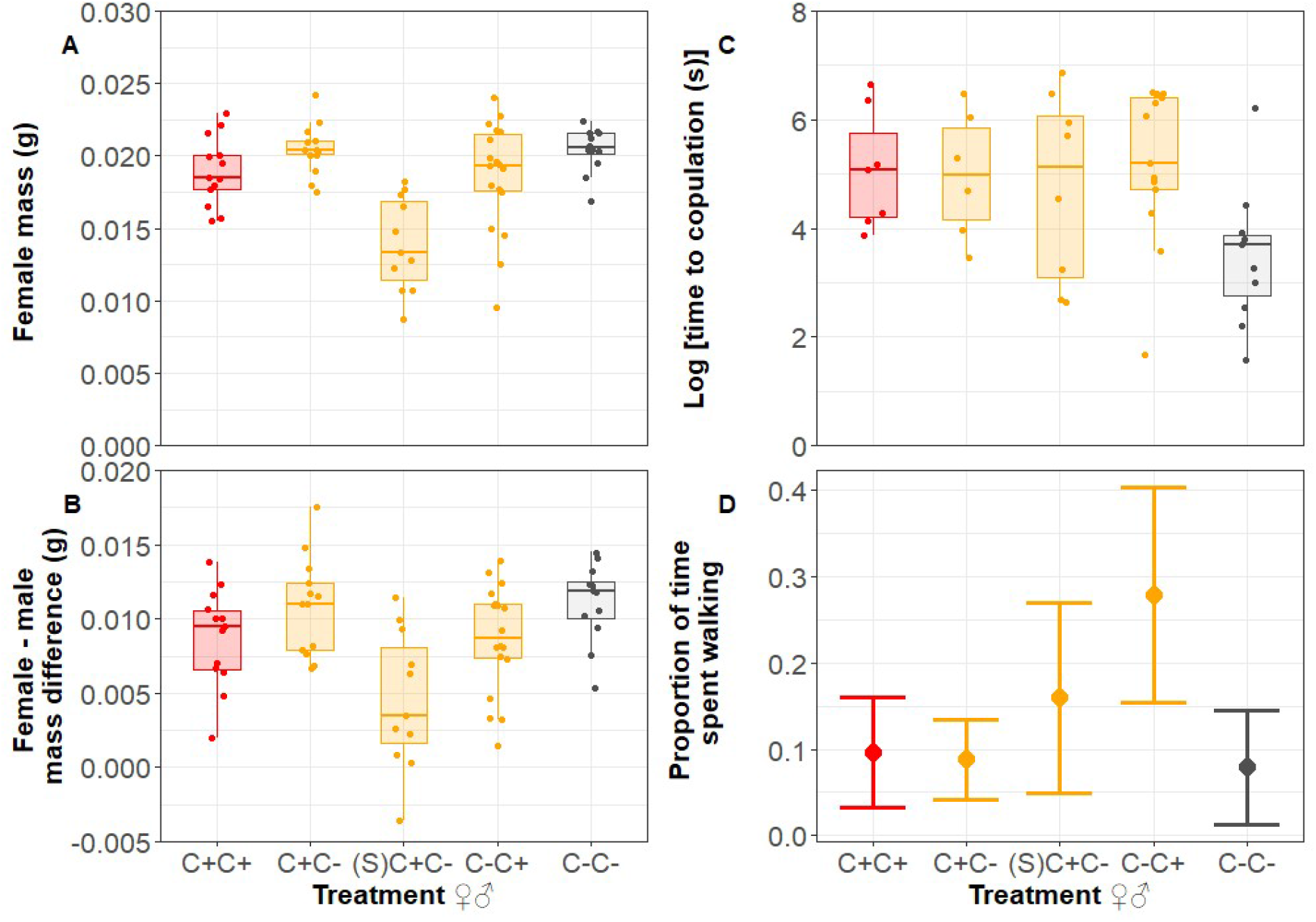
Effect of clerodanoid access (C+ = nibbled on *Ajuga reptans* leaf, C- = not nibbled on *A. reptans* leaf) and early life starvation of females [(S)C+] on A) body mass, B) difference in body mass between adult females of *Athalia rosae* and the males they were paired with, C) time until copulation, and D) proportion of time spent walking during copulation (given as estimate marginal mean). Treatment levels are listed with females on the left and males on the right. Boxes in box plots show the median, the first and third quartiles (the 25th and 75th percentiles) at the hinge, and the whiskers extend to the largest or smallest value no further than 1.5 * IQR from the hinge for the upper and lower whiskers, respectively.

There was no effect of treatment on the occurrence (X^2^_4,62_ = 7.72, p = 0.102) or the rate of leg battling (X^2^_4,62_ = 4.53, p = 0.339) during the assays. Treatment also did not significantly affect the occurrence of fighting/attempted nibbling (X^2^_4,62_ = 6.68, p = 0.154), its rate (X^2^_4,62_ = 8.11, p = 0.088), or its duration (Pseudo-R^2^ = 0.131, (LRT) X^2^ = 6.36, df = 4, p = 0.174). Both the occurrence and rate of successful nibbling events did vary significantly with treatment (occurrence: X^2^_4,62_ = 14.13, p = 0.007; rate: X^2^_4,62_ = 13.36, p = 0.010), being higher in pairs with asymmetrical clerodanoid access (figures 3A & 3B), however this difference was only significantly different for nibbling rate in (S)♀C+♂C- and ♀C-♂C+ treatment levels when compared to the control (table 2). This successful nibbling on conspecifics was also predominantly driven by C-individuals (X^2^_1,66_=31.81, p<0.001, figure 4). In contrast there was no effect of treatment on nibbling duration (Pseudo-R^2^ = 0.068, (LRT) X^2^ = 1.24, df = 4, p = 0.870). The likelihood of having an agonistic interaction pre-copulation was significantly affected by treatment (X^2^_4,40_ = 11.00, p = 0.026, figure 3C). Time until copulation commenced was also longer in pairs where pre-copulatory fighting occurred (X^2^_1,43_ = 11.08, p =0.002, figure 3D). An interruption of copulation by agonistic behaviour was common across all treatment levels (X^2^_4,40_ = 8.92, p = 0.063), and did not significantly influence copulation duration (X^2^_1,43_ = 1.27, p= 0.266).

**Figure 3.**
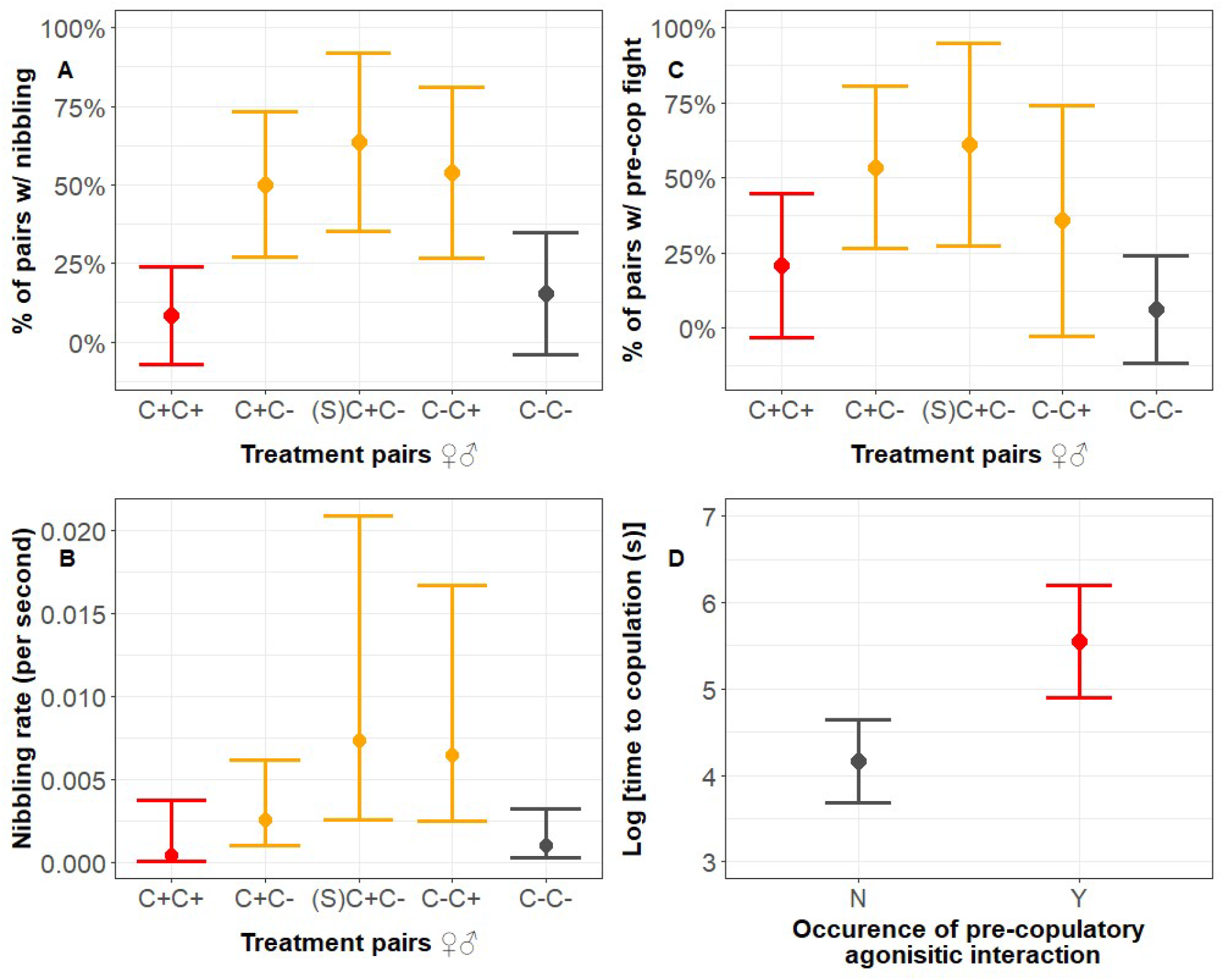
Effect of clerodanoid access (C+ = nibbled on *Ajuga reptans* leaf, C- = not nibbled on *A. reptans* leaf) and early life starvation of females [(S)C+] on A) the occurrence of successful nibbling, B) the rate (or frequency) of successful nibbling events, C) agonistic interactions prior to copulation, and D) time taken until copulation commence depending on whether pre-copulatory agonistic behaviour also occurred in adult *Athalia rosae*. Treatment levels/pairs are listed with females on the left and males on the right. Plots show model predictions with 95% confidence intervals.

**Figure 4.**
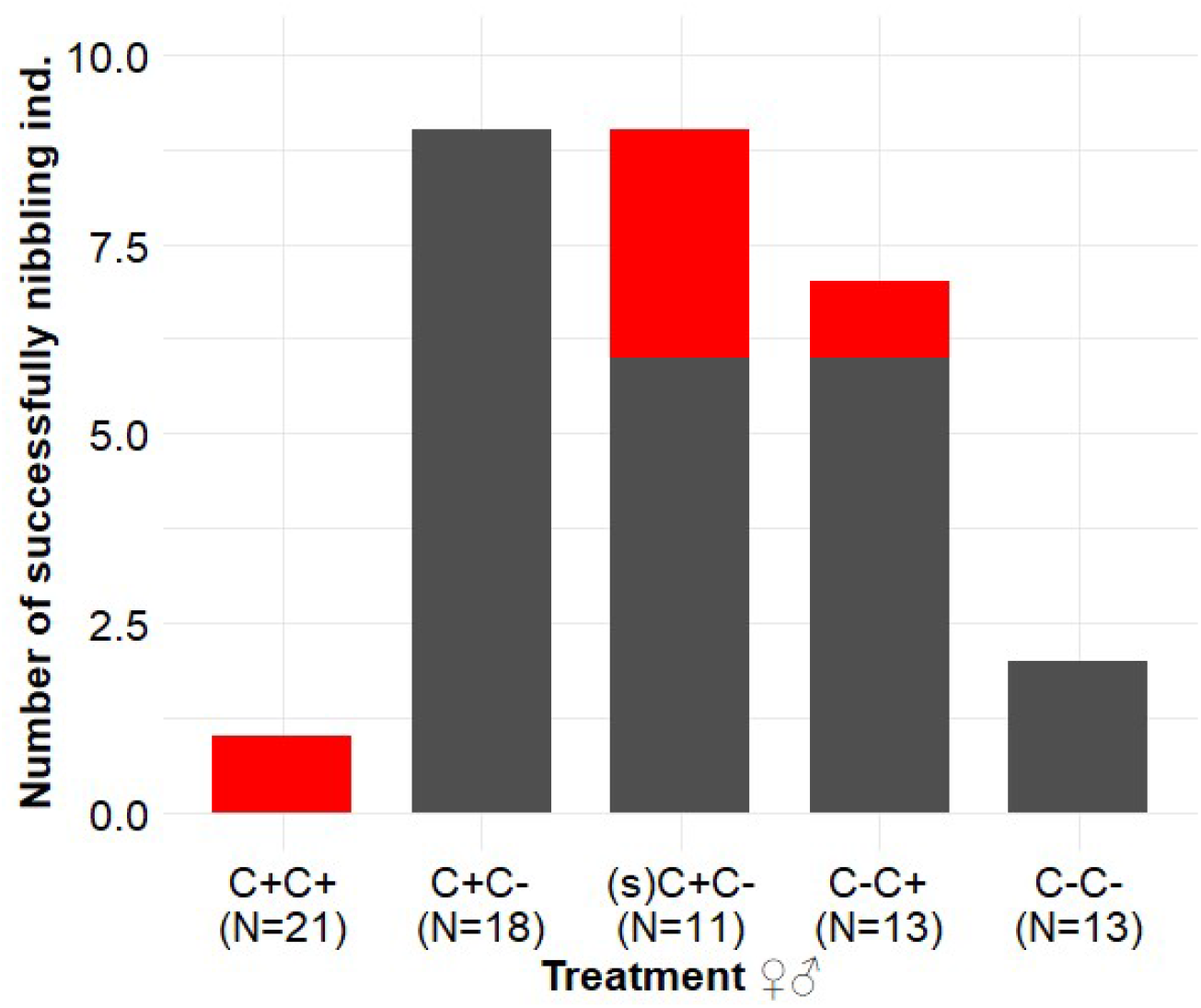
The number of adult *Athalia rosae* individuals that successfully nibbled their opponent by treatment, split within treatment by whether or not they had clerodanoid access (**red**; C+ = nibbled on *Ajuga reptans* leaf) or not (**grey**; C- = not nibbled on *A. reptans* leaf). Total number of pairs in each treatment level (N), given on x axis.

**Table 2.** Results of pairwise comparisons (Tukey HSD) between different treatment levels assessing the drivers of the significant main effect of treatment on the occurrence and rate of successful nibbling behaviour in adult *Athalia rosae*. C+ = clerodanoid access (nibbled on *Ajuga reptans* leaf), C- = no clerodanoid access (not nibbled on *A. reptans* leaf), and (S) = starvation of females during larval development.

| Treatment | Nibbling occurrence |  |  |  | Nibbling rate |  |  |  |
| --- | --- | --- | --- | --- | --- | --- | --- | --- |
|  | Estimate | Std. Error | z | p | Estimate | Std. Error | z | p |
| ♀C+♂C+ vs ♀C-♂C- | -6.93E-01 | 1.30E+00 | -0.53 | 0.946 | -8.50E-01 | 1.21E+00 | -0.70 | 0.892 |
| ♀C+♂C- vs ♀C-♂C- | 1.70E+00 | 9.02E-01 | 1.89 | 0.169 | 9.06E-01 | 7.24E-01 | 1.25 | 0.531 |
| **(S)♀C+♂C-vs ♀C-♂C-** | **2.26E+00** | **9.92E-01** | **2.28** | **0.070** | **1.97E+00** | **7.75E-01** | **2.54** | **0.038** |
| ♀C-♂C+ vs ♀C-♂C- | 1.86E+00 | 9.49E-01 | 1.96 | 0.147 | **1.84E+00** | **7.43E-01** | **2.48** | **0.046** |

## Discussion

The main aim of this study was to investigate the degree to which access to clerodanoids influences both mating and fighting behaviour in *A. rosae*. Here we show that agonistic behaviour commonly occurs between male and female *A.rosae* and that it can delay the onset of copulation, although it does not prevent copulation from occurring. In line with our predictions, compared to the control where neither individual had clerodanoids, the acquisition of clerodanoids by females had a stronger positive effect on mating behaviour than when males had clerodanoids, although this was not the case for all mating parameters. Also as predicted, successful nibbling increased in asymmetrical clerodanoid treatments compared to the control and was highest when RHP differences between attacker (C-) and defender (C+) were in the attacker’s favour. As we discuss in detail below, these findings support the idea that fighting between intersex pairs over clerodanoids plays a key role in determining mating behaviour in *A. rosae*, further highlighting the importance of studying intersex agonistic interactions.

In general, agonistic behaviour between individuals was high across all treatment levels, potentially indicating a degree of sexual conflict independent of clerodanoids, which warrants further investigation. Such agonistic behaviour may be expressed to assess a partner’s quality or rather to avoid costly copulations [50], especially in haplodiploid species like *A. rosae* where unmated females can still produce male offspring [41]. Pairs of *A. rosae* with pre-copulatory fighting took significantly longer to commence copulation, with the occurrence of pre-copulatory agonistic behaviour being higher in the two treatment levels with C+ females and C-males (♀C+♂C- and (S)♀C+♂C-). This difference in the occurrence of pre-copulatory fighting may go some way to explaining the weak trend for the commencement of mating in asymmetrical treatment levels (♀C+♂C-, (S)♀C+♂C-,♀C-♂C+) compared to the control (♀C-♂C-) being longer than the commencement of mating in the symmetrical C+ treatment level (♀C+♂C+) compared to the control. The occurrence of precopulatory fighting in these asymmetrical pairs also suggests that despite the potential benefits of mating with a C+ female, males are willing to forgo immediate mating in order to gain clerodanoids. Whether such agonistic behaviour in the wild would mean a lost mating opportunity is difficult to fully assess here as the interactions occurred in an enclosed arena, which in turn may contribute to overall high mating success across treatments. It does, however, demonstrate the importance of acquiring clerodanoids to males i.e. high RV. This is in spite of the fact that such acquisition did not seem to positively influence male mating success; for example, when paired with a C-female C+ males had a higher rate of failed copulation attempts in comparison to control pairs.

Despite generally high levels of agonistic behaviour across treatment levels, only (S)♀C+♂C- and ♀C-♂C+ showed an increased rate of successful nibbling. That this should be the case arguably reflects differences in body mass, and therefore RHP [51], between the individuals with or without clerodanoids in these two asymmetrical treatment levels. In *A. rosae* females are larger and heavier than males [38,39] and therefore in ♀C-♂C+ contests the C-individual (female) is more likely to be able to overpower the C+ individual (male), whereas the reverse is true in ♀C+♂C-contests. Along similar lines in (S)♀C+♂C- the C+ females have reduced mass meaning that the C-males are also more likely to be able to gain clerodnaoids from them in a contest compared to normal sized females. That body mass is a key determinant of the direction and success of fights and nibbling interactions over clerodanoids in *A. rosae* fits well with numerous studies on size and RHP in other species [52–55] and highlights the importance of sexual dimorphism in determining the direction of intersex contest interactions. The enhanced rate of successful nibbling behaviour in ♀C-♂C+ pairs may also go some way to explaining the higher number of failed copulations and female walking during copulation observed in this treatment level compared to the control, reflecting the female’s unwillingness to mate opposed to nibble. One might have expected a similar effect on mating parameters in the (S)♀C+♂C-treatment, but this was not the case. However, there was a difference between the two treatments in the sex which had and did not have clerodanoids, which indicates that for attacking C-males it is still beneficial to mate with a C+ female, even when competitor RHP is reduced.

Female quality effects could also have fed into results seen in the (S)C+C- treatment level. Early life effects such as starvation can negatively affect an individual’s fighting ability independent of size effects [56]. Therefore (S)C+ females may have had a stronger reduction in RHP than perhaps would be expected merely from the lower body mass of these females compared to well-fed individuals, increasing the likelihood of successful nibbling by males. Furthermore, the effect of early life starvation on females may have shifted the balance between the costs and benefits of fighting before mating (or vice versa) via a decrease in female quality. Early life stressors such as starvation are known to affect sexually selected traits [57] in a way that has negative effects on mating and reproductive success [58–60]. Starved *A. rosae* females were lighter and size is strongly linked to reproductive success in many insects [61]; thus, (S)C+ females may have been less attractive as mating partners. Conversely, early life starvation can also result in poor quality females that are less choosy [62], making mating more likely in pairs from a female choice perspective. Here it is difficult to disentangle the various effects of early life starvation on RHP and mate quality, but future experiments could do so via matching for size between sexes irrespective of early life treatment [e.g. 63].

In addition to providing anti-predator defence [33] previous work, showing that males mated more quickly with C+ females but that females showed no preference for C+ males, has also suggested that clerodanoids may act as a female pheromone [36]. Through their effect on female survival clerodanoids could arguably provide a reliable binary signal of female quality and therefore be good candidates for compounds that act as pheromones. Clerodanoid uptake provides *A. rosae* individuals with better protection against some predators [33] resulting in C+ females being potentially longer lived than C-females, due to their lower predation risk, and subsequently laying a larger number of eggs (increasing fitness). Furthermore, such a dual role of clerodnaoids would fit the general pattern of infochemical flexibility observed in insects [64,65] and other animals [66]. The results for mating behaviour presented here weakly support the notion that C+ females have increased mating success whereas C+ males do not, and therefore the idea that clerodanoids may act as a pheromone [35]. However, we also clearly demonstrate how the disruption of mating behaviour by fighting over clerodanoids can contribute to such differences in mating success and unlike the previous study we did not use a mate choice assay, so such support is far from conclusive. Niether are both explanations - mate attraction vs. fighting over defence compounds – mutually exclusive, both could be contributing to the behaviour observed in this experiment. Additional measures of both male and female investment in response to mate C+ status e.g. sperm [67,68] or egg [69,70] quality and quantity measures, may help to further unravel this puzzle by giving us an indication of how clerodanoids influence perceived mate quality and therefore their role as potential pheromones.

In summary, we investigated agonistic intersex interactions over a resource between potential mating partners and to our knowledge show for the first time that fighting over such a resource can have knock-on effects on mating behaviour, particularly the onset of mating. The success of aggressive nibbling behaviour in pairs with asymmetrical clerodanoid exposure was determined by the superiority of RHP in C-individuals. Poor mating success of C+ males, i.e. the number of failed mating attempts, seems to have been driven by the agonistic nibbling behaviour of C-females, which had a higher RHP than males. What is more, the reduction of the RHP of C+ females via early life starvation increased successful agonistic nibbling behaviour in pairs with a C-male. Further work is needed to disentangle the effects of early life stress on factors determining fighting ability and mate quality and the effect this has on male-female interactions such as the ones outlined here, as well as to generally assess the roles of RHP, RV and pre-fight assessment in determining interaction occurrence and duration in this species. Overall, we demonstrate the importance of studying agonistic intersex interactions and hope that this work stimulates further research in the area.

## Supporting information

S1 a - Fighting video

S1 b - Mating video

## Acknowledgements

Many thanks to M Höning for assistance maintaining the *A. rosae* cultures and S Lane for feedback on an earlier version of the manuscript. This research was funded by the German Research Foundation (DFG: https://www.dfg.de/) as part of the SFB TRR 212 (NC^3^) – Project number 396777467 (granted to CM).

## Author contributions

CM conceived the study. SP designed the study and carried out experimental rearing, behavioural assays, coding of behavioural data, statistical analysis, and wrote the manuscript. CM helped to significantly revise the manuscript. Both authors gave final approval for publication.

## Ethics statement

Research was carried out following the Association for the Study of Animal Behaviour / Animal Behavior Society Guidelines for the Use of Animals in Research (Animal Behaviour, 2020, 159, I-XI) and following guidance laid out in Drinkwater, Eleanor, Elva JH Robinson, and Adam G. Hart. “Keeping invertebrate research ethical in a landscape of shifting public opinion.” Methods in Ecology and Evolution 10.8 (2019): 1265-1273. Individuals were collected in the field using a sweep net to catch them in flight or when sitting on flowers, eliminating excess bycatch (approx < 0.1% of catch). After the behavioural assays individuals were returned to the *A. rosae* stock culture.

## Data Availability

Data (.csv files) and annotated scripts (R) used in analyses are available from Zenodo repository (https://doi.org/10.5281/zenodo.4537747).

