## Supplementary material for "Fighting over defence chemicals disrupts mating behaviour": S2

S2. Results of pairwise comparisons (Tukey HSD) between different treatment levels assessing the drivers of the significant main effect of treatment on both female mass (g) and female-male mass difference (g).

| **Treatment** | **Female mass (g)** | | | | | **Female-male mass difference (g)** | | | |
| --- | --- | --- | --- | --- | --- | --- | --- | --- | --- |
|  | **Estimate** | **Std. Error** | **z** | **p** | | **Estimate** | **Std. Error** | **z** | **p** |
| ♀C+♂C+ vs ♀C+♂C- | 1.80E-03 | 1.03E-03 | 1.74 | | 0.406 | 2.44E-03 | 1.32E-03 | 1.85 | 0.342 |
| ♀C+♂C+ vs ♀C-♂C+ | -5.13E-06 | 1.11E-03 | -0.01 | | 1.000 | 2.91E-04 | 1.41E-03 | 0.21 | 1.000 |
| ♀C+♂C+ vs ♀C-♂C- | 1.49E-03 | 1.11E-03 | 1.34 | | 0.664 | 2.31E-03 | 1.41E-03 | 1.64 | 0.473 |
| ♀C+♂C- vs ♀C-♂C+ | -1.81E-03 | 1.01E-03 | -1.79 | | 0.378 | -2.15E-03 | 1.29E-03 | -1.67 | 0.451 |
| ♀C+♂C- vs ♀C-♂C- | -3.13E-04 | 1.01E-03 | -0.31 | | 0.998 | -1.25E-04 | 1.29E-03 | -0.10 | 1.000 |
| ♀C-♂C+ vs ♀C-♂C- | 1.49E-03 | 1.09E-03 | 1.37 | | 0.643 | 2.02E-03 | 1.39E-03 | 1.46 | 0.588 |
| **(s)♀C+♂C- vs ♀C+♂C+** | **6.52E-03** | **1.16E-03** | **-5.65** | | **<0.001** | **6.56E-03** | **1.47E-03** | **-4.45** | **<0.001** |
| **(s)♀C+♂C- vs ♀C+♂C-** | **4.72E-03** | **1.06E-03** | **-4.46** | | **<0.001** | **4.12E-03** | **1.35E-03** | **-3.05** | **0.020** |
| **(s)♀C+♂C- vs ♀C-♂C+** | **-6.53E-03** | **1.13E-03** | **-5.76** | | **<0.001** | **-6.27E-03** | **1.45E-03** | **-4.33** | **<0.001** |
| **(s)♀C+♂C- vs ♀C-♂C-** | **-5.04E-03** | **1.13E-03** | **-4.44** | | **<0.001** | **-4.24E-03** | **1.45E-03** | **-2.93** | **0.028** |
